# Generation of Pathway Signatures by Combining Normal Modes with Weighted Ensemble Simulations

**DOI:** 10.64898/2026.04.21.719877

**Authors:** Anthony T. Bogetti, Anupam Banerjee, Ken Dill, Ivet Bahar

## Abstract

Molecular dynamics simulations provide a “computational microscope” by which molecular phenomena can be studied at atomic resolution. However, such simulations are often expensive, usually due to a combination of system size and timescale. Various enhanced sampling methods have been proposed to overcome these challenges. Despite their effectiveness, many suffer from artifacts from energetic biases guiding the simulations, or lack of effective progress coordinates. Proteins’ normal modes uniquely defined by their 3D fold capture their intrinsic dynamics and could provide unbiased guidance, but how to combine these modes with molecular dynamics to generate continuous, energetically unbiased pathways has been challenging. In this study, we demonstrate that conformations generated along from normal modes using adaptive anisotropic network model provide a physical, intuitive, and generalizable progress coordinate for weighted ensemble simulations, providing a boost in efficiency and a means to generate pathways for any protein system without prior knowledge.

## Introduction

As is often the case, there is no “absolute best” method for simulating biomolecules like proteins. Often, there is a sharp tradeoff between model detail and the time cost of the simulated data. Take for instance, molecular dynamics (MD) simulations, which are atomically detailed, and which often contain explicit water molecules and ions; these simulations can be very expensive and limited to relatively short time scales or relatively small sized systems, but are often rewarding, as the resulting data are also high in detail. On the other hand, simple, analytical models like elastic network models (ENMs)^1-4^ are very quick to compute and provide rich information on a protein’s intrinsic dynamics, and especially on the large-scale cooperative movements of supramolecular systems,^5,6^ but at a coarse-grained resolution.

Efforts to make MD simulations more efficient often rely on coarse-graining^7,8^ or employing enhanced sampling methods.^9-12^ In either case, there are challenges and tradeoffs, but often the results are more useful and insightful than those standalone conventional MD simulations could provide. Enhanced sampling methodologies generally fall into two categories: methods that employ energetic biases, or those that apply sampling biases.^13^ Either category relies on one or more progress coordinates along which a “bias” is applied. The weighted ensemble (WE) strategy is such an enhanced sampling method that focuses MD on barrier crossings, “cutting out” the waiting time spent in the stable states.^14-16^ It relies heavily on a carefully-chosen progress coordinate to enhance sampling. Once an appropriate coordinate is found, WE can generate energetically unbiased, continuous pathways, enabling the exploration of mechanisms of a wide variety of processes such as chemical reactions,^17^ protein folding,^18-20^ drug permeation through a membrane,^21^ protein-protein binding,^22,23^ protein-ligand unbinding,^24-26^ and protein conformational change.^27,28^ However, the efficiency gains of WE simulations are often countered by the challenges of defining an effective progress coordinate, and extensive knowledge of the system of interest is often a prerequisite. Thus, the promise of the WE strategy is great, but the reality is often limited by lack of effective progress coordinate.

Efforts to make the results from ENMs more detailed often rely on building multi-step protocols, or attempting to combine elastic network models with MD. These methods, called hybrid methods,^29,30^ can take on many forms depending on the question or system.^31-35^ One particular hybrid method, the adaptive Anisotropic Network Model (aANM),^32^ proceeds through iterative rounds of model building and structural deformation, with the goal of exploring the most probable, or energetically most favorable, pathways between known endpoints. While the ANM is based on a simplified model of structure and energetics (an elastic network with uniform spring constants), the predicted residue fluctuations correlate with NMR and X-ray B-data^36^; and more importantly, the soft modes of motions obtained by ANM normal mode analysis have been shown to be highly robust^4,37^, and relevant to cooperative functional events otherwise observed in atomic simulations of micro-to-milliseconds^38^. ANM pathways are fast to compute but are coarse-grained. This means that analysis of detailed interactions formed or broken during the pathway is often challenging. Thus, again, the promise of aANM is great, but the reality is often limited by the lack of detail in the resulting pathway.

Combining methods can often result in highly effective “super methods,”^39,40^ but sometimes the details of *how* to combine the methods can be tricky to figure out, though recent attempts have proven the utility of simulating specific types of conformational change processes using WE and a single soft mode as a progress coordinate.^41^ That being said, a combined WE-aANM method could be highly effective, providing WE with a coarse-grained pathway intrinsically favored by the network architecture, which it could use as a progress coordinate; and aANM offers such a pathway that can be translated into fully atomic, high-resolution trajectories (**Figure 1**). In this work, we develop a method that combines WE with aANM for maximum effect, named AWE-PATH: <u>A</u>NM-guided <u>WE</u> generation of <u>PATH</u>S. Our method, in theory, builds on the string method protocol for WE simulations, and is carefully implemented in the widely used software packages WESTPA 2.0^42^ and *ProDy* 2.0.^43^ Because this protocol is based on WE, users can make use of the rich set of tools available for WE analysis, e.g., directly analyze ensembles for extracting kinetics data without having to rely on external analysis suites. We illustrate the application of AWE-PATH to many systems of varying size and complexity demonstrating how the combination of these two powerful methods, rather than any one method, provides an effective means of sampling transitions at high resolution with no prior knowledge of the behavior of the proteins being simulated.

**Figure 1.**
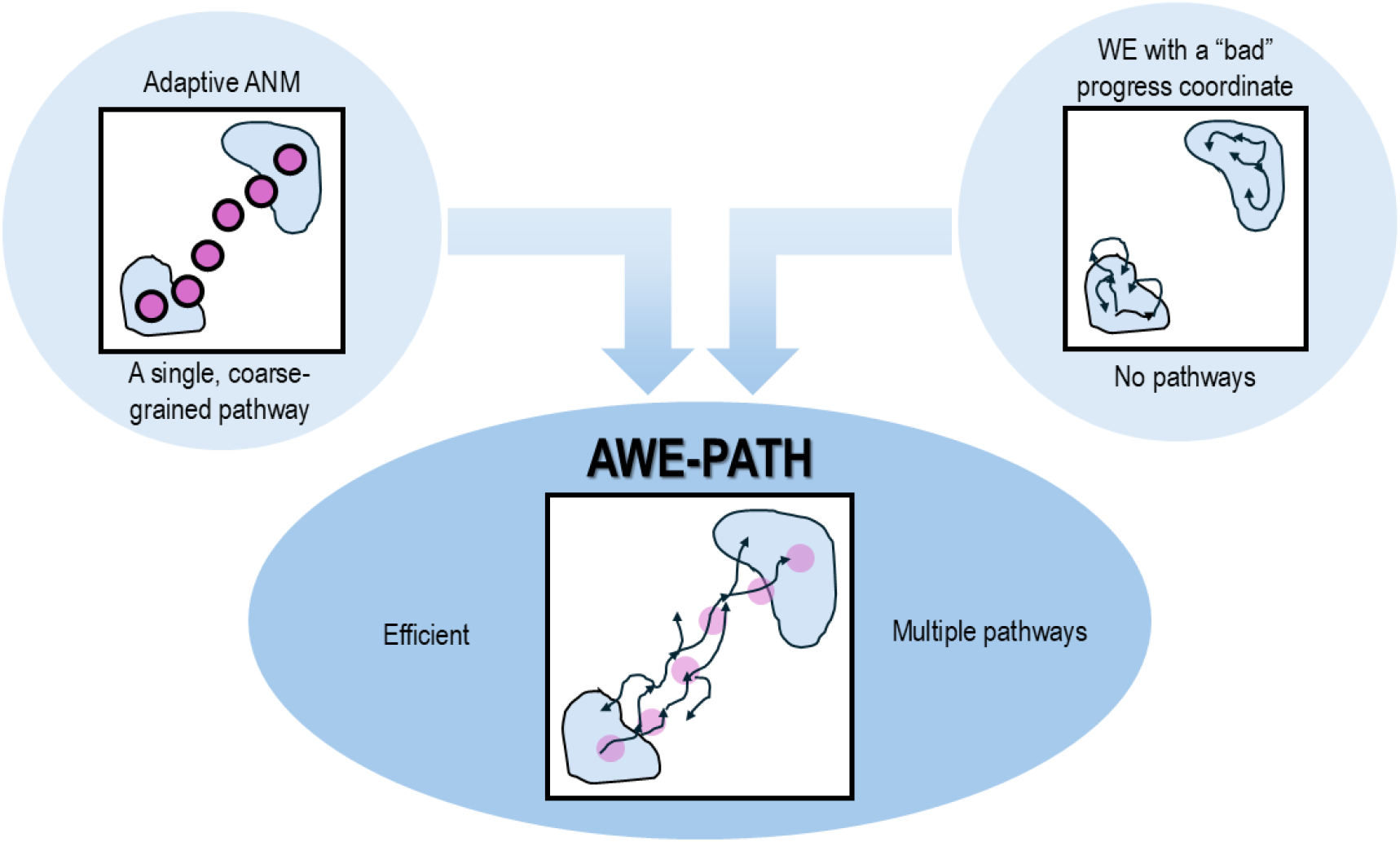
AWE-PATH combines adaptive ANM and WE simulations for maximum effect. Pathways from adaptive ANM provide general and efficient progress coordinates for WE simulations. While on their own, each method can generate pathways, the combination of adaptive ANM (aANM) and weighted ensemble (WE) into a “super method” can generate atomically detailed pathways along collective motions uniquely defined by the architecture in a generalizable and efficient manner.

## Results

### AWE-PATH: A new tool for ANM-guided WE simulations of transition mechanisms

We present below the major ingredients of AWE-PATH: (i) a variant of the adaptive ANM (aANM) method that provides an initial coarse-grained pathway or a series of transient conformers (*milestones*) along the softest collective modes intrinsically accessible to the endpoints; (ii) a new algorithm that uses these conformers to define a progress coordinate for generating an ensemble of WE trajectories.

### Adaptive ANM (aANM) variant for guiding AWE-PATH

aANM has been proposed for efficient sampling of conformational transitions undergone by supramolecular systems between two endpoints A and B^32^. The transition is effectuated via a succession of normal modes adaptively reevaluated as the system, represented by an ANM, moves away from its original conformation, thus yielding a ‘coarse-grained’ pathway. We adopted here a variant of aANM to adapt to WE. The protocol begins with constructing the ANMs^44^ for the energy-minimized structures A and B, i.e., the 3*N*-dimensional vectors composed of the 3D position coordinates of *N* α-carbons, designated as ***R***_***A0***_ and ***R***_***B0***_. These are iteratively updated to generate a series of *transient* (or *intermediate*) conformers, starting from both ends. For generating the first transient conformer ***R***_***A1***_, we determine a subset of low frequency (also called *soft*) modes of motions accessible to ***R***_***A0***_, by ANM normal mode analysis. Note that these modes robustly entail cooperative movements of large substructures/domains often relevant to function,^4,6,37,45^ well beyond conventional MD. ***R***_***A0***_ is deformed along a subset of soft modes that yields a sufficient overlap with the initial 3*N*-dimensional deformation vector ***d***_*0*_ = ***R***_***B0***_ - ***R***_***A0***_. Here, sufficient overlap refers to a cumulative squared overlap criterion, whereby the minimal subset of low-frequency modes is selected such that their combined contribution exceeds a prescribed threshold, ensuring that the deformation remains aligned with the target direction while favoring intrinsically soft collective motions. The magnitude of each deformation step is further scaled to limit the displacement along the selected modes and maintain the validity of the harmonic approximation. Different choices of overlap threshold and step-size scaling may lead to alternative, yet physically plausible, pathways connecting the same endpoints; full algorithmic details are provided in the Supplementary Methods.

The resulting new transient conformer, ***R***_***A1***_, serves as input for the next cycle. The procedure is repeated at the opposite end ***R***_***B0***_ as well, to adaptively generate the transient conformer ***R***_***B1***_, and the new distance vector ***d***_***1***_. This back-and-forth iterative process, termed an aANM cycle, is terminated at the *k*^*th*^ cycle, provided that there is a “meeting” in the middle defined by an RMSD of <1.5Å between ***R***_***Ak***_ and ***R***_***Bk***._ The number of intermediates, *M*, is bound to an upper limit (i.e., 40).

The aANM method thus adaptively produces a series of coarse-grained conformers at C^α^ resolution representing milestones along an easily accessible (soft) pathway between the endpoints, albeit at low resolution. Note that a multitude of pathways along different combinations of normal modes may achieve the transition. However, the goal is to find the minimal path lengths that encounter the lowest energy barriers via optimal selection of aANM parameters (see Supplementary Methods for details). The aANM pathway is then used in WE simulations to define a progress coordinate. All aANM pathways were generated with the aANM module added to the open source application programming interface (API) *ProDy 2*.*0*.^43^

### WE algorithm and its implementation in AWE-PATH

The WE algorithm is an efficient, highly scalable, and statistically exact method used in conjunction with MD simulations for enhanced sampling of conformational space, including rare events and transitions, without altering energetics or adding forces that bias the underlying dynamics.^46^ Each WE iteration consists of an MD productive run for a fixed time interval, τ, followed by resampling during which the trajectories are extended, split, or merged depending on the subspace (bins) sampled, to achieve an adequate coverage of the conformation space. The probabilities (or weights) of the ensemble of trajectories emanating from a parent trajectory sum up to unity at all times. WE shares characteristics with a subgroup of enhanced sampling methods, broadly called path sampling strategies, like transition path sampling,^47^ transition interface sampling,^48^ milestoning^49^ and forward flux sampling.^50^ By rigorously assigning and tracking trajectory weights, it has a benefit over other path sampling strategies in that rates can be directly computed from the generated pathways through analysis of fluxes (probability flow over time).^46^

The AWE-PATH framework, depicted in **Figure 2**, takes an aANM pathway, and converts it into an efficient and interpretable 1D coordinate for guiding WE simulations. To this aim, the aANM pathway (**Figure 2a**) is first generated and then projected onto a 2D RMSD space (**Figure 2b**), the *x*- and *y*-axes of which represent the RMSD in the coordinates of the respective conformations A and B with respect to their initial energy-minimized structures ***R***_***A0***_ and ***R***_***B0***_. This space is general and will describe any two-state transition. The aANM pathway is embedded into this 2D RMSD space as a polyline consisting of segments linking the end points A and B via intermediate conformers (C) along the aANM pathway (magenta polyline in **Figure 2b**). Next, a second polyline is constructed using the midpoints of conformations from the magenta polyline, giving the cyan polyline in **Figure 2b**.

**Table 1.** WE simulations using an aANM-based progress coordinate are faster in generating pathways than their standard radially binned counterparts. In case of our showcase systems, the speedup was often 2 to 3 times as fast and is expected to become more efficient as the system size and complexity increases.

| Protein | WE resampling type | Transition A → B |  | Transition B → A |  |  |
| --- | --- | --- | --- | --- | --- | --- |
| | | Time to 1<sup>st</sup> pathway (μs) | Speedup | Time to 1<sup>st</sup> pathway (μs) | Fraction of productive runs ($f$) | Speedup ($f$) |
| Adenylate kinase | Radial bins | 0.21 +/- 0.02 | 1.62 | No pathways | 0/3 | ∞ (2/3) |
|  | aANM path | 0.13 +/- 0.06 |  | 0.20 +/- 0.01 | 2/3 |  |
| Ribose-binding protein | Radial bins | 0.12 +/- 0.07 | 2.00 | 0.15 +/- 0.00 | 1/3 | 1.88 (1/3) |
| | aANM path | 0.06 +/- 0.01 | | 0.08 +/- 0.02 | 3/3 | ∞ (2/3) |
| Maltose-binding protein | Radial bins | 0.38 +/- 0.39 | 3.80 | 0.43 +/- 0.25 | 2/3 | 1.54 (2/3) |
| | aANM path | 0.10 +/- 0.02 | | 0.28 +/- 0.14 | 3/3 | ∞ (1/3) |

**Figure 2.**
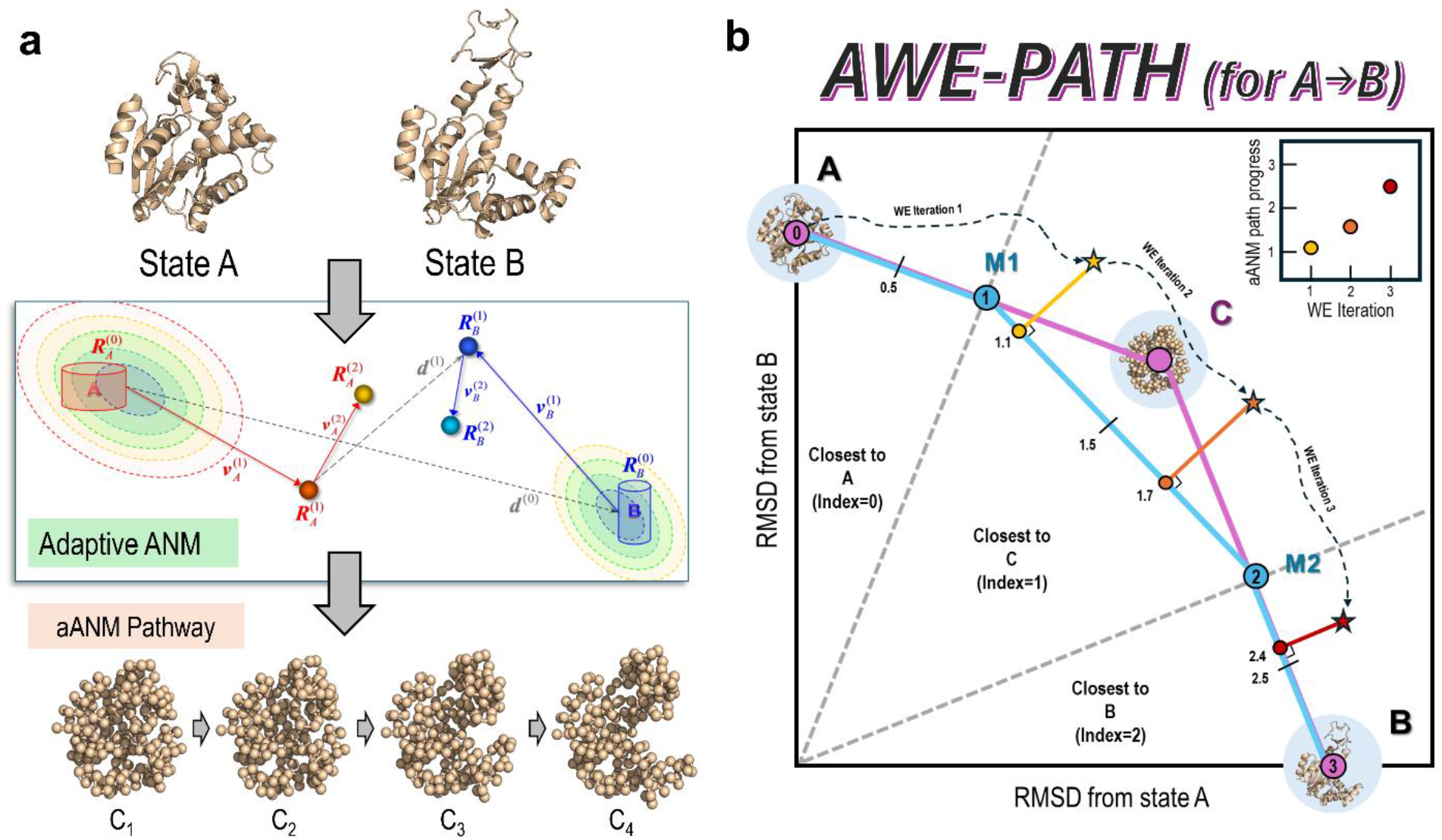
The AWE-PATH framework compresses an aANM pathway into a 1D progress coordinate by way of a 2D polyline. **(A)** The first step of AWE-PATH is to generate a pathway with adaptive ANM, which provides between end-point conformations A and B, *M* transient conformations (*M* = 1 in this schematic representation), labeled as C. **(B)** Next, the adaptive ANM pathway, including the end points, is projected onto a 2D RMSD space. A polyline is then formed based on the midpoints between successive conformations in the aANM pathway, with the number of line segments in the polyline equal to *M* + 2. During the AWE-PATH simulation’s resampling stage, the instantaneous WE conformations (shown by the *yellow, orange and red stars*) are evaluated with respect to this polyline to identify which polyline segment they are closest to. In this example, the first instantaneous WE conformation (*yellow star*) is closest to the segment connecting M_1_ and M_2_. Thus, the instantaneous progress would be 1 + a fractional value that represents how close the next line segment is. In this example, the *yellow star* has a progress of 1.1 because it is only 10% of the way to the next line segment (M_2_-B).

For each instantaneous WE conformation (yellow, orange and red stars in **Figure 2b**), we calculate the RMSD to A and RMSD to B, locating that conformation in the 2D RMSD space. The RMSD calculations are performed with Amber’s cpptraj program.^51^ Then, the position of the WE conformation along the cyan polyline is evaluated as: progress= *X* +*F*, where *X* is an integer and refers to the segment that is closest to the WE conformation (0 ≤ *X*˂*M*), and *F* accounts for the fractional progress along that particular segment (0 ˂*F* ˂1). The AWE-PATH framework thus effectively evaluates progress along the inherently high dimensional aANM pathway in a low-dimensional way, letting the WE algorithm “interpret” the structural information from the aANM pathway in a flexible manner. The measure of progress is one dimensional and can be used in WE like any other progress coordinate. The actual code implementation of AWE-PATH is explained further, with pseudo-code and details, in the Supplementary Methods. For binning, AWE-PATH uses the minimal adaptive binning (MAB) scheme^52^ to adaptively generate bins as our simulations progress. Along the 1D progress coordinate, five direction-specific MAB bins and one direction-specific bottleneck bin were used along with a target number of five trajectories per bin.

*Conventional WE as a control*. The results from AWE-PATH were compared to those from conventional WE simulations. The conventional WE runs were set up in the most general and efficient way for pathway generation, using the above 2D RMSDs as a 2D progress coordinate.^53^ Radial binning was adopted, with bins placed every 5° from 0° to 90°, as schematically depicted in **Figure S1**.This type of partitioning of the conformational space allows for relatively efficient transit between A and B without too many unproductive splitting in regions of high A or B RMSD, like a grid-based binning scheme would promote.

### AWE-PATH generates accurate pathways for three medium-sized proteins

We first present the results for three test proteins, adenylate kinase (AK) (**Figure 3a**), ribose-binding protein (RBP) (**Figure 3b**), and maltose-binding protein (MBP) (**Figure 3c**). All three proteins are widely studied/characterized proteins, which permit us to better interpret the results in terms of prior work, as well as control WE simulations. **Table S1** lists the PDB structures used as endpoints for each protein, the corresponding protein size in number of residues, and RMSDs between endpoints. In every case, we assigned the more compact structure as A and the less compact structure as B. We performed six independent AWE-PATH simulations (three A → B and three B → A) of 150 WE iterations each, with time interval τ of 100 ps; and we combined the generated trajectories (with conformers saved at 10 ps intervals) into a single datafile for each protein. The combined trajectories were then projected onto the 2D RMSD subspace. Projections of the same simulation data onto physical coordinates relevant to each protein (i.e., the nmp and lid opening angles for AK) are presented for reference in **Figure S2**. The selected physical coordinates for the three respective proteins, as plotted in the Supplemental Information, are: (i) the angle between the core and nucleoside monophosphate (NMP) domains (*x*-axis) and the angle between the Core and Lid (ATP-binding) domains (*y*-axis), consistent with previous work^31,54-56^ (ii) the hinge and twisting angle between the two domains of RBP,^57-59^ and (iii) the radius of gyration and domain opening angle of MBP.^60-62^ In each of the above-mentioned physical spaces, the aANM conformers and resolved structures specific to the protein pair are also projected for reference. Data from conventional WE simulations using the radial binning scheme depicted in **Figure S1** are similarly plotted in the 2D RMSD subspace (**Figure S3**) and the physical subspaces described above (**Figure S4**).

**Figure 3.**
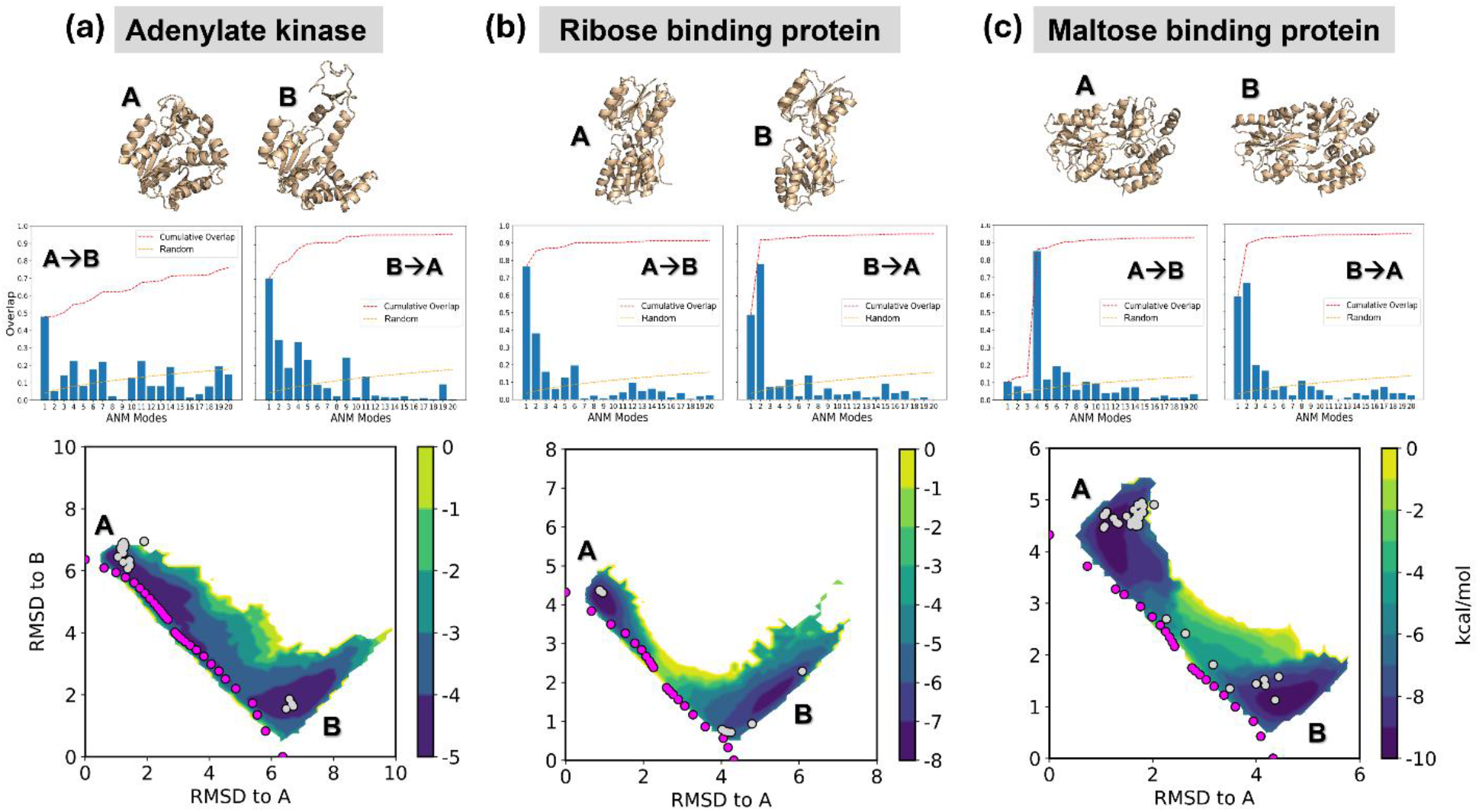
AWE-PATH simulations broadly sample the conformational space between end point conformations for three medium-sized showcase protein systems, (a) adenylate kinase, (b) ribose-binding protein, and (c) maltose-binding protein. Each distribution shown is the combined result of six independent WE simulations, three for the A to B direction and three for the B to A direction, with WE weights normalized across the six independent simulations using the w_multi_west tool. Magenta dots represent the location of the aANM pathway and grey dots represent resolved structures for each protein with the same uniport ID. See Table S1 for more details on the three systems. The mode spectra for the deformation of the A→B and B→A transitions, calculated from ANMs of the end point structures A and B, demonstrate that the transitions are enabled by the soft modes.

AWE-PATH simulations yield an ensemble of pathways for each system, which were either broadly distributed (as in AK, see **Figure S2**) or narrowly confined to the transition direction (RBP and MBP) (**Figures S3 and S4**). Two major differences between AWE-PATH and conventional WE were: (a) the forward passages (from open to closed states) were achieved more easily by AWE-PATH compared to control WE, as indicated by the shorter time to first pathway listed in **Table 1**; and (b) the reverse transition (from closed to open), which is more challenging in general, could not be achieved in the majority (6/9 runs) of control WE simulations; whereas AWE-PATH failed to reach the end point in 1/9 case only. The speedup achieved by the aANM-driven WE compared to conventional WE during the forward transition was observed to be larger (up to ∼4 times), as the system size and complexity increases. Our new protocol is expected to be particularly useful for large systems, consistent with the distinguished efficiency of ANM for modeling the collective dynamics of supramolecular systems.

Comparison with earlier studies, in addition to that with conventional WE with radial binning, lends support to AWE-PATH results. For example, previous simulations revealed a multiplicity of pathways for the transition of AK between its open and closed states, which is consistent with the heat map in **Figure S2a**.^56^ Notably, LID closure seems the first event during the transition from open to closed state (see magenta dots in **Figure S2a**), consistent with previous simulations^56,63,64^ in the apo state; whereas, starting from the closed state, viable paths are the opening of either NMP or LID domains (along the vertical and horizontal axes), or their coupled movements (along the diagonal). The holo state was reported to exhibit such interdomain coupling, i.e., AMP binding that facilitate the closure of the NMP domain, would in turn, stabilize the closed state of the LID^55^. Notably, the respective time scales of the closed → open and open → closed transitions of apo AK were reported by Zheng and Cui to be 1.8 - 77.2 μs and 12.2 - 357.7 μs^56^, in qualitative accord with the relative time scales (or occurrences) of these two passages observed here. Haran and coworkers observed opening rates of ∼45 microseconds by smFRET^56,65^.

In contrast to AK, RBP and MBP exhibit relatively well-defined transition pathways (**Figure S2b** and **c**). Previous study has shown that the transition of RBK between its open and closed states can be accounted for by three softest ANM modes describing the twisting, bending and rotational motions of its two domains with respect to each other,^56,59^ hence the success of ANM-guided WE in both directions. The timescale of PDB conformational change has been reported to be on the fast ns timescale.^57^ As to MBP, which undergoes a “Venus-flytrap mechanism,” the motion is similar to that of RBP, a similarly simple transformation, but with a detectable intermediate “semi-open” structure.^61^ The timescale of closing was reported to be on the ns to μs timescale by Tang et al. from paramagnetic relaxation experiment NMR.^66^

### AWE-PATH generates accurate pathways for two proteins selected from a database

We performed a real-world test applying AWE-PATH to generate transition pathways for two proteins’ endpoints selected from the CoDNaS 2.0 database or protein conformational change.^67^ For each protein system, we ran a single A→B and a single B→A AWE-PATH simulation and then combined the resulting distributions much like we did with the showcase systems presented earlier. The resulting distributions, combining two WE simulations each, one A→B and one B→A, were normalized and plotted in the 2D RMSD space, along with the aANM conformers (magenta dots) and resolved structures of the same protein (grey dots).

AWE-PATH was able to successfully sample transitions for both proteins, as shown in **Figure 4**. The first protein was ribonuclease J, which is 150 residues in size and undergoes a conformational change of around 9 Å RMSD. Our AWE-PATH simulations were successfully able to sample the entire conformational space between the two end points (**Figure 4a**), generating a continuous pathway connecting the two end states in as little as 0.17 μs of aggregate sampling time. Ribonuclease J conformational change here involves docking and undocking of a terminal helix, which is easier in the B to A direction as indicated by the mode spectra and AWE-PATH simulations.

**Figure 4.**
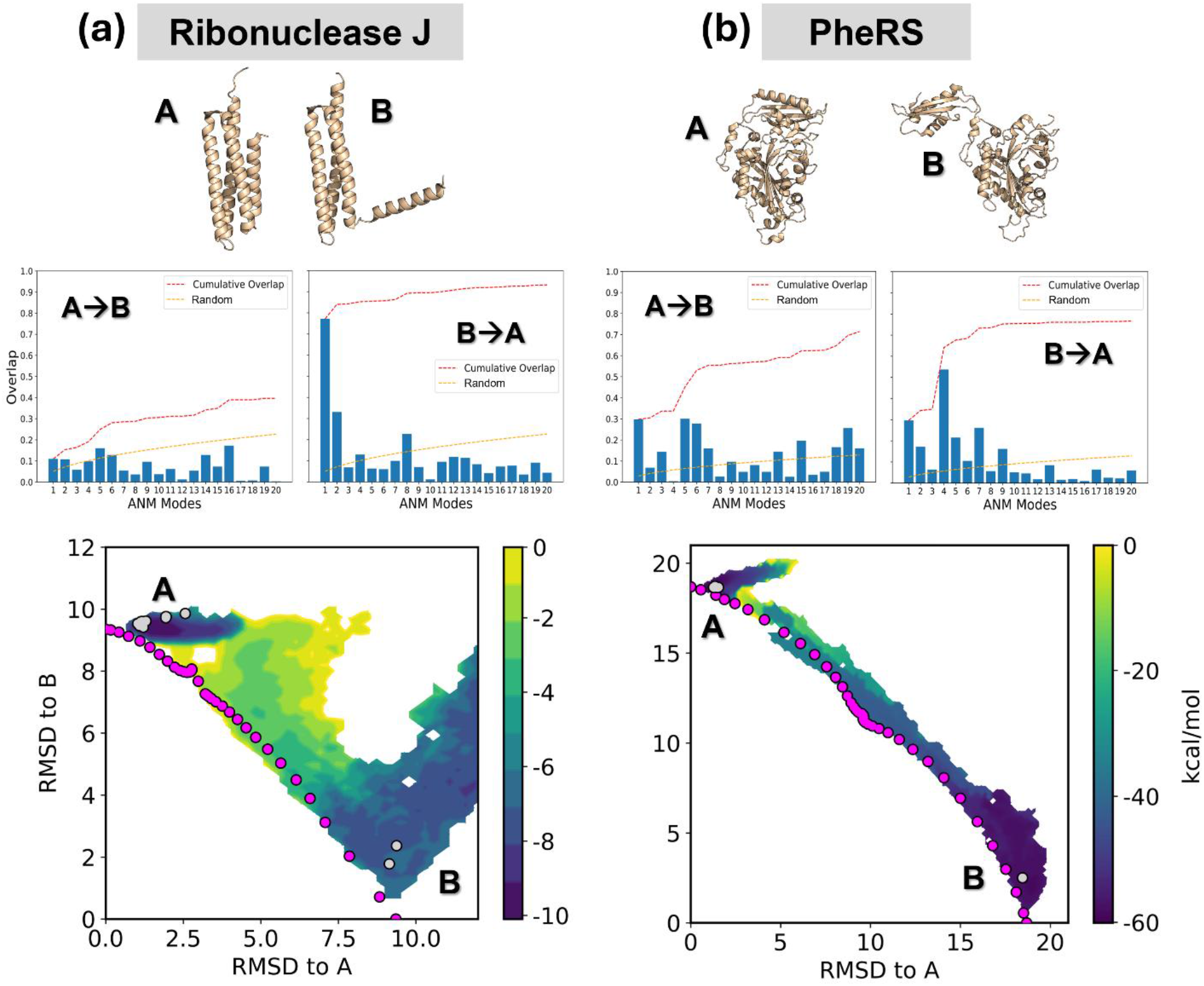
AWE-PATH sampled complete transitions for two proteins selected from the CoDNaS 2.0 database, (a) ribonuclease J and (b) PheRS. The distributions shown are the combined result of two independent WE simulations, one for the A to B direction and one for the B to A direction, with WE weights normalized across the two independent simulations. Magenta dots represent the location of the aANM pathway and grey dots represent resolved structures for each protein. See Table S1 for more details on the systems. The mode spectra for deformations in the A→B and B→A transitions demonstrate that the transitions are enabled by the soft modes in at least one of the two transition directions.

The second protein, PheRS, is 404 residues in size and undergoes a conformational change of nearly 20 Å RMSD, making this system the largest and most complex conformational change simulated so far. Nevertheless, AWE-PATH was able to fully sample the conformational space connecting the two end states (**Figure 4b**) and generated a continuous pathway in only 0.15 μs of aggregate sampling time. The motion for PheRS conformational change involves the rotation of a domain around a flexible linker. The mode spectra for this conformational change indicates that the motion is enabled by the softest modes.

### AWE-PATH generates accurate pathways for a stress-test protein

As a final stress-test for AWE-PATH, we applied it to the large-scale conformational change involving the opening/closing of HIV-1 reverse transcriptase (**Figure 5**). This system represents a stress test for our method due to its large size of 959 residues. AWE-PATH was able to sample the conformational space between end points in only 1.2 μs of aggregate simulation time. In addition, resolved structures of HIV-1 reverse transcriptase (grey dots in **Figure 5**) overlap with key regions of sampled space from AWE-PATH simulations, including a transition region between A and B, indicating the accuracy of AWE-PATH in sampling physical transitions.

**Figure 5.**
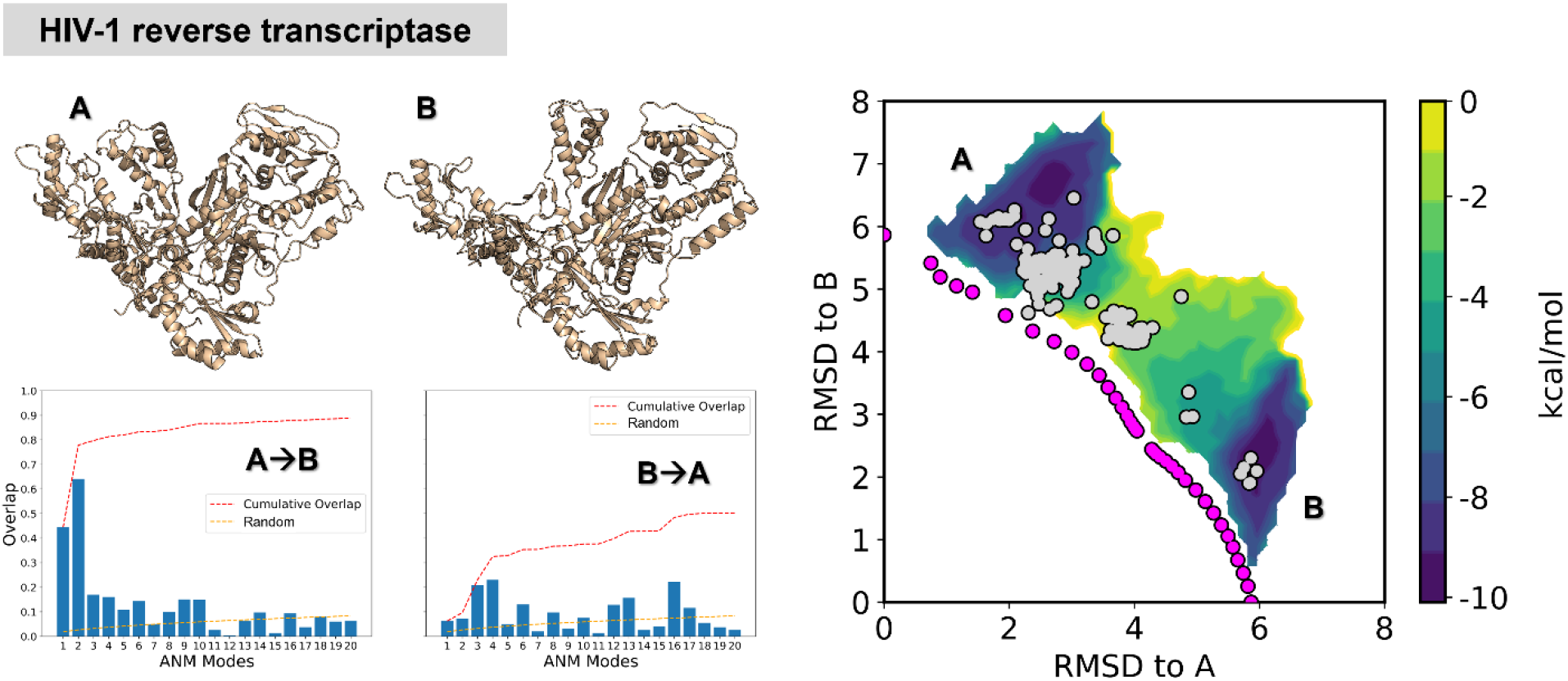
AWE-PATH sampled complete transitions for HIV-1 reverse transcriptase. The distribution shown is the combined result of two independent WE simulations, one for the A to B direction and one for the B to A direction, with WE weights normalized across the two independent simulations. Magenta dots represent the location of the aANM pathway and grey dots represent resolved structures for each protein. See Table S1 for more details on the systems. The mode spectra for each direction demonstrate that the A→B and B→A transitions are enabled by the soft modes in at least one of the transition directions.

## Conclusions

Energetically unbiased pathways provide a clear view into the mechanism of protein action. However, generating these pathways is not always trivial, as path sampling strategies often rely on user-defined metrics of progress that can be hard to rationalize. While machine learning has helped in the identification of such generalizable coordinates, there is still a way to go to generate coordinates that are both effective and physically understandable. In this study, we have demonstrated that an initial pathway from aANM can serve as a physical sampling guide for WE, and we developed the AWE-PATH framework in which an aANM pathway can be described as a low-dimensional progress coordinate for use in WE simulations. We generated transition pathways for three showcase proteins, two proteins selected from the CoDNaS 2.0 database and a stress-test protein with AWE-PATH, and through comparing the sampled space with the location of resolved structures, confirmed the accuracy of AWE-PATH in generating physically meaningful pathways in an efficient manner.

## Acknowledgements

IB gratefully acknowledges support from NIH awards R01 GM139297-06 and R01 DA062680 and from WoodNext Foundation.

