## Supplemental Information for "Generation of Pathway Signatures by Combining Normal Modes with Weighted Ensemble Simulations"

### Supplemental Methods

**Adaptive Anisotropic Network Model.** Adaptive anisotropic network model (aANM) analysis was used to generate coarse-grained conformational transition pathways between two endpoint structures, $A$ and $B$. The method constructs pathways by iteratively deforming each structure along collective low-frequency normal modes that are recalculated at every step, allowing the deformation directions to adapt to changes in contact topology as the structure moves away from its initial conformation.

The endpoint conformations are represented as 3N-dimensional coordinate vectors

$$R_A^{(0)}, R_B^{(0)}$$

where $N$ is the number of $C_\alpha$ atoms. At iteration $k$, the instantaneous deformation vector is defined as

$$d_k = R_B^{(k)} - R_A^{(k)}$$

An anisotropic network model is constructed for the current structure, and its non-rigid-body normal modes $\left\{u_i^{(k)}\right\}$ are obtained from diagonalization of the ANM Hessian.

To determine which modes contribute to the deformation at a given iteration, the overlap of each mode with the instantaneous deformation vector is computed as

$$O_i^2 = \frac{(u_i^{(k)}.d_k)^2}{\|d_k\|^2}$$

Modes are selected cumulatively, starting from the lowest-frequency end of the spectrum, until the sum of squared overlaps satisfies

$$\sum_{i \in S_k} O_i^2 \geq F_{min}$$

where $S_k$ denotes the selected subset of modes at iteration $k$. This criterion ensures that the deformation direction remains sufficiently aligned with the target direction while favoring intrinsically soft, collective motions. At least one mode is always selected. When not specified explicitly, $F_{min}$ is determined adaptively and capped at an upper bound (0.6 in this work) to limit excessive recruitment of higher-frequency modes.

The deformation direction is constructed as a weighted linear combination of the selected modes,

$$v_k = \sum_{i \in S_k} (u_i^{(k)}.d_k) u_i^{(k)}$$

The structure is then updated according to

$$R_A^{(k+1)} = R_A^{(k)} + s_k v_k$$

The step size $s_k$ is chosen to optimally reduce the instantaneous distance to the target along the selected deformation direction and is given by

$$s_k = f\frac{v_k . d_k}{v_k . v_k}$$

where the unscaled factor corresponds to the optimal projection of the deformation vector onto the selected mode subspace. The scaling factor $f$ controls the magnitude of the displacement and limits each step to remain within the regime where the harmonic approximation underlying normal mode analysis is valid. Smaller values of $f$ result in more conservative, incremental deformations, whereas larger values accelerate convergence at the risk of unphysical distortions. In this work, a value of $f = 0.2$ was used to balance numerical stability and computational efficiency.

An analogous procedure is applied to the structure originating from the opposite endpoint, yielding a back-and-forth iterative process in which both structures move toward one another. This alternating construction reduces directional bias and allows the pathway to reflect collective motions intrinsic to both endpoint conformations. The iterative process is terminated when one or more convergence criteria are met: (i) the RMSD between the two evolving structures falls below a target threshold (1.5 Å), (ii) successive RMSD reductions become negligible, or (iii) unphysical deformations such as loss of network connectivity are detected. The total number of intermediate conformers generated along the pathway is limited to a maximum of 20.

Because the deformation at each step depends on the selected modes and scaling parameters, the aANM framework does not produce a unique transition pathway. Different choices of the overlap threshold $F_{min}$, step-size scaling factor $f$, and limits on the number of modes can lead to alternative pathways that nonetheless connect the same endpoints. Lower $F_{min}$ values favor motion along fewer, softer modes and typically yield longer, lower-energy excursions, whereas higher $F_{min}$ values recruit additional modes and produce more direct pathways. The parameter values used here were chosen to balance physical plausibility, numerical stability, and suitability for integration with weighted ensemble sampling.

**The AWE-PATH algorithm.** The AWE-PATH algorithm operated in 2D space with the reference aANM pathway as a polyline. The algorithm is direction-specific; it can operate in either the A to B direction or the B to A direction. In this discussion, we will present an example demonstrating the algorithm applied to the A to B direction. The reference aANM pathway polyline has multiple segments connecting end point conformations A and B. If the aANM pathway contained a single transient conformation C, then the polyline would be composed of two segments, $\overline{AC}$ and $\overline{CB}$. Now, for the purposes of our algorithm, we generate a second polyline composed of the segment midpoints from the first polyline. For example, our previous polyline of A-C-B would be used to generate A-$M_{\overline{AC}}$-$M_{\overline{CB}}$-B, where $M_{\overline{AC}}$ is the midpoint along the line segment $\overline{AC}$ and $M_{\overline{CB}}$ is the midpoint along $\overline{CB}$. We will rename these as $M_1$ and $M_2$, respectively, giving us the following polyline for use in our progress coordinate calculation: A-$M_1$-$M_2$-B. This gives us a total of three line segments, $\overline{AM_1}$, $\overline{M_1M_2}$ and $\overline{M_2B}$.

Next, for each WE conformation W, we determine its position along the AWE-PATH coordinate according to the following algorithm. The big picture is to find (i) which segment W is "closest" to, and (ii)) where does the projection of W fall onto that closest segment. The answer to the first question provides an integer value that is greater than 0 but less than 3. This value corresponds to which conformation, A (0), C (1) or B (2), W is closest to in the 2D RMSD space. For instance, if W is closest to $\overline{AM_1}$, then it is closest to index 0, or conformation A. Only after it surpasses $M_1$ does it become most similar to C, with an index of 1. Passing $M_2$ indicates W would be most similar to B with an index of 2, which is the highest integer possible because B is the end of the original A-C-B polyline. Thus, the largest possible value for the AWE-PATH progress coordinate is *M*+1 where *M* is the total number of conformations from aANM. One may notice that this part of the algorithm is placing W inside a Voronoi bin created from the original A-C-B polyline.

The answer to the second question will be a decimal value and will indicate how far W has to go before it changes which segment it is most similar to. Say that W is “closest” to the $\overline{AM_1}$ segment of the A-$M_1$-$M_2$-B polyline. It’s index is 0, which means it is closest to conformation A, but how close is it to become most similar to C? Remember that passing $M_1$ would mean that W would be closest to C, so we quantify how far W traveled along the $\overline{AM_1}$ segment as a decimal value between 0 and 1. If W is exactly halfway, it would be 0.5 of the way to C, for example. This decimal value allows AWE-PATH to finely locate W along the polyline and further quantify progress along A-C-B in a better way than simple Voronoi bins could.

Here is the pseudocode for the algorithm, where a point W and midpoint list M (i.e. [$M_1$,$M_2$ ...]) are used as inputs and the output is list of floating point values that falls within the range 0 to len(M)+1. This algorithm is run for each WE trajectory T in the current iteration, with the trajectory T composed of multiple WE conformations W.

```
1. LOAD the "current" WE trajectory (T).

2. LOAD the "midpoint" list (M) that defines the path.

3. FOR each point W in T:

   a. INITIALIZE the "minimum distance" as infinity.

   b. FOR each pair of consecutive midpoints in the path (M[i] as M1 and M[i+1] as M2)

      i. CREATE VECTORS:

         - Vector_M1M2 = M2 - M1 (The direction of the segment)

         - Vector_M1W = W - M1  (Point W relative to the segment start)

      ii. CALCULATE PROJECTION:

         - Find the squared length of the segment: LengthSq = (Vector_M1M2 · Vector_M1M2).

         - IF LengthSq is 0:

            - The "closest point" is simply M1.

         - ELSE:

            - Find the relative position: Ratio = (Vector_M1W · Vector_M1M2) / LengthSq.

            - CLAMP the Ratio between 0.0 and 1.0 to stay on the segment.

            - Calculate the "closest point" = M1 + (Ratio * Vector_M1M2)

      iii. CALCULATE “CLOSENESS”:

         - Find the squared distance between W and the "closest point".

      iv. UPDATE BEST MATCH:

         - IF this squared distance is the smallest seen so far for point W:

            - Store the current segment index (i) as the "Base Component".
```

- Store the clamped Ratio as the "Bonus Component".

c. OUTPUT FINAL PCOORD VALUE: Base Component + Bonus Component (e.g., 1.25).

The CLAMP step ensures that line segments do not extend beyond their boundaries and generate negative numbers for the Bonus Component. In addition, the “IF LengthSq is 0” check remedies situations where two subsequent midpoints (M[i] and M[i+1]) may actually be the same point, which sometimes occurs if the adaptive ANM conformations barely changed from one round of deformation to the next (note that if our recommendation to take a subset of the aANM pathway is followed, the likelihood of this happening is very low, and even if it does happen this check will ensure the score is still properly calculated).

**System preparation for simulations.** Protein Data Bank[1] **(**PDB) structure files were downloaded and prepared with the pdb4amber program.[2] Missing internal loops were built using Modeller.[3] Each protein was assigned force field parameters from the FF19SB force field,[4] solvated in a truncated octahedral box of explicit OPC waters, with sodium or chloride Li-Mertz 12-6-4 ions added to achieve neutrality. The padding between the protein and the edge of the box varied between system and conformation, with a larger padding for more compact systems (so that when they opened, there would be sufficient clearance with the edge of the box). Each system was then energy minimized for 2,000 steps. For all subsequent MD, the pmemd.cuda engine as part of the Amber20 software package[5] was used. A timestep of 2 fs was used and periodic boundary conditions were applied to the system with particle mesh Ewald electrostatics and a long-range cutoff of 10 Å. The energy minimized structure was heated to 300 K over the course of 20 ps in an NVT ensemble with weak (1 kcal/mol) positional restraints on all protein heavy atoms. A Langevin thermostat was used with a friction coefficient of 1 per ps. Following heating to 300 K, the system was simulated with the same positional restraints for 1 ns in an NPT ensemble with a Monte Carlo barostat with a coupling constant of 1 per ps. In the final equilibration, the positional restraints were released, and the system was simulated for another 1 ns in an NPT ensemble.

**WE trajectory analysis.** For each protein, we ran simulations of 150 iterations each in the A $\rightarrow$ B direction, and, separately, in the B $\rightarrow$ A direction. In each direction, once trajectories fell below 1.5 Å RMSD to the target conformation (see rectangular bins in **Figure 2**), they were recycled, i.e., the trajectory was terminated and its weight was transferred to a trajectory back in the starting state. Recycling is a technique commonly used to accelerate convergence to a steady state probability distribution in WE simulations. We used a dynamics interval of 100 ps for all systems and saved intermediate structures for analysis at 10 ps intervals. All WE simulations in this study were run using the open source WESTPA 2.0 software package.[6]

We combined the generated trajectories (or intermediates saved at 10 ps intervals) into a single datafile using the w_multi_west tool. We plotted the resulting normalized average probability distribution of all conformations sampled by WE with their associated weights using the WEDAP plotting tool.[7] Conditional probability fluxes and event duration times were calculated with the built-in WESTPA 2.0 analysis tools w_assign and w_direct using states for A and B as defined in **Figure 2**. Pathway signatures were generated by extracting successful trajectories (those that reached the target state of A or B, depending on the simulation direction) using the built-in w_trace tool.

#### Supplemental Tables

**Table S1. Structures whose transitions were simulated by AWE-PATH.**

| Protein name | PDB code for closed | PDB code for open | Number of residues | Starting $C^{\alpha}$ RMSD (Å) |
|---|---|---|---|---|
| Adenylate kinase | 1AKE[8] | 4AKE[9] | 214 | 6.36 |
| Ribose binding protein | 2DRI[10] | 1URP[11] | 271 | 4.32 |
| Maltose binding protein | 1ANF[12] | 1OMP[13] | 370 | 4.32 |
| Ribonuclease J | 7W9U | 7W8B | 150 | 9.35 |
| PheRS | 5MGH[14] | 3TUP[15] | 404 | 18.68 |
| HIV-1 reverse transcriptase | 1HQE[16] | 1VRT[17] | 959 | 5.87 |

#### Supplemental Figures

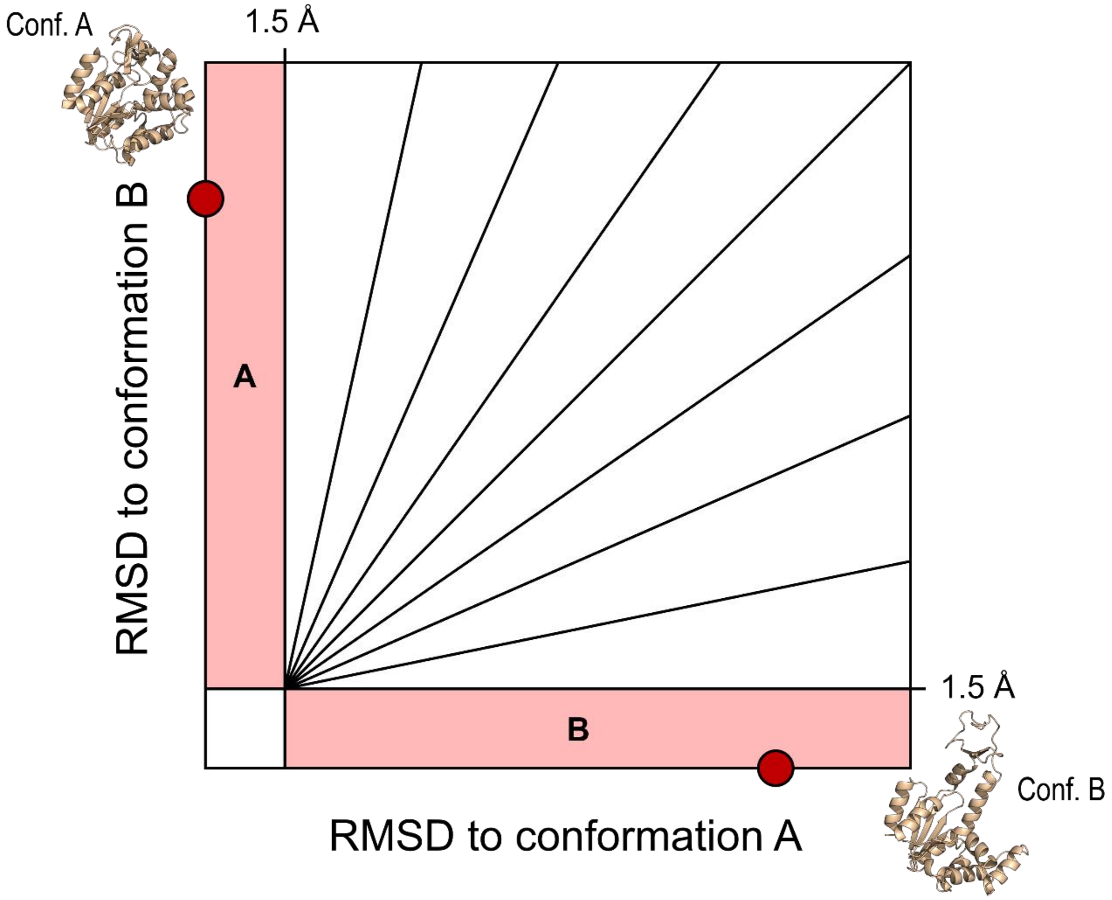


**Figure S1. A radial binning scheme for conventional WE simulations**. Our conventional WE simulations employed a 2D progress coordinate. The first dimension of this coordinate was the $C^{\alpha}$ RMSD to conformation A and the second dimension the $C^{\alpha}$ RMSD to conformation B. Below 1.5 Å in each dimension was considered that conformation's stable state and emanating out from (1.5,1.5) were radial bins that were placed every 5° from 0° to 90°. The diagram above shows fewer radial bins than were used during our simulations, for illustrative purposes.

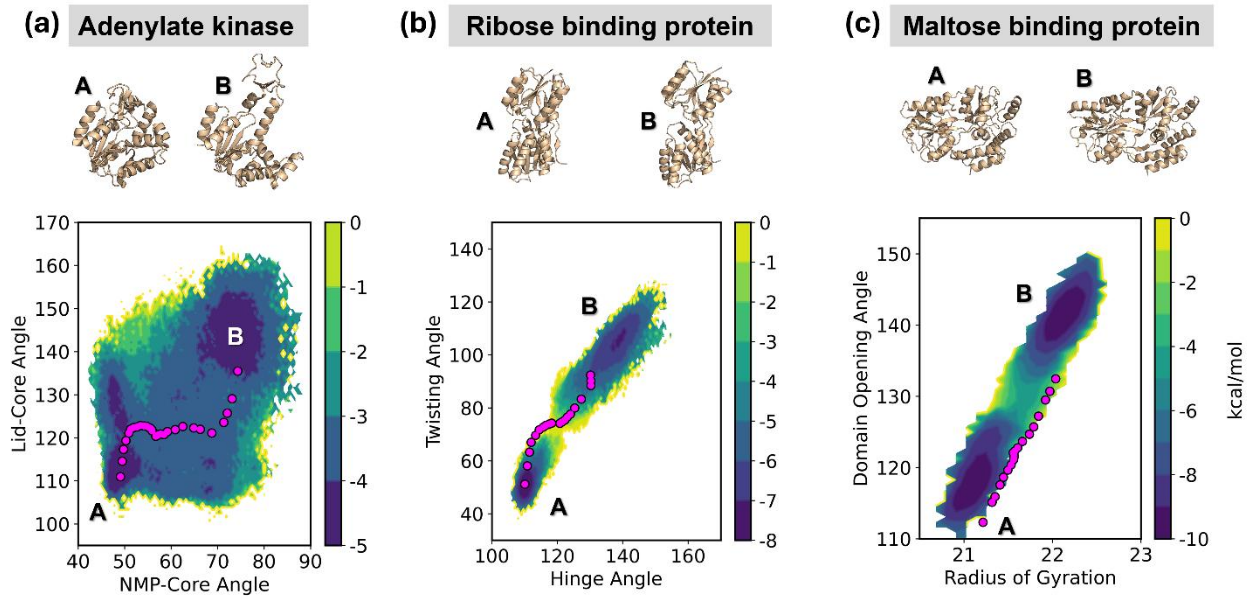


**Figure S2. AWE-PATH simulations plotted in common, physical coordinate subspaces for three medium-sized showcase protein systems, (a) adenylate kinase, (b) ribose-binding protein, and (c) maltose-binding protein.** The distributions shown are the combined result of six independent WE simulations, three for the A to B direction and three for the B to A direction, with WE weights normalized across the six independent simulations. Magenta dots are the aANM pathway conformers.

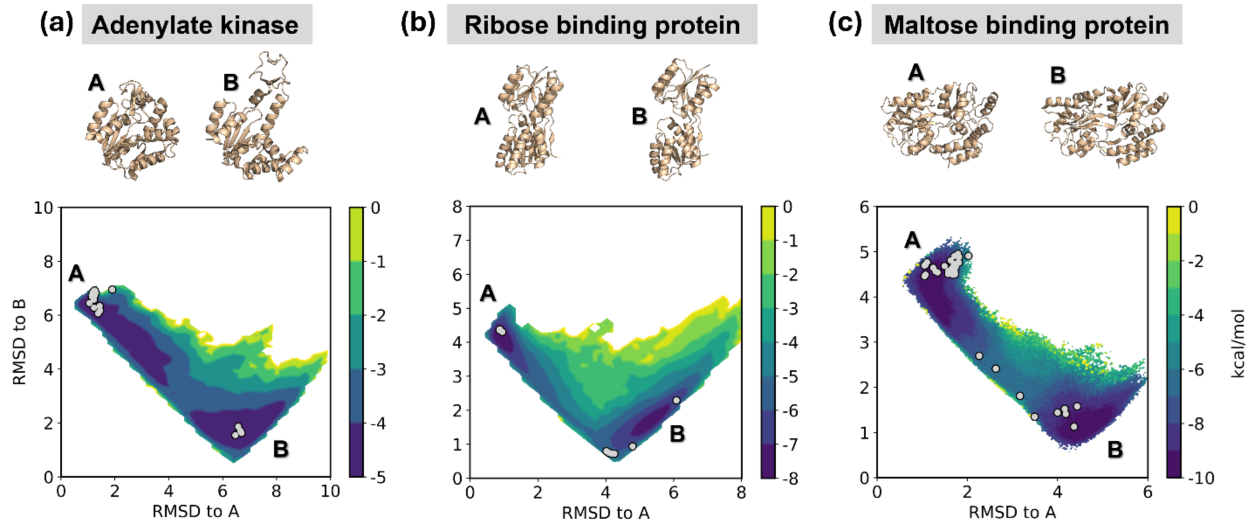


**Figure S3. Conventional WE simulations with a radial binning scheme broadly sample the conformational space between end point conformations for three medium-sized showcase protein systems, (a) adenylate kinase, (b) ribose-binding protein, and (c) maltose-binding protein.** The distributions shown are the combined result of six independent WE simulations, three for the A to B direction and three for the B to A direction, with WE weights normalized across the six independent simulations. Grey dots represent resolved structures for each protein.

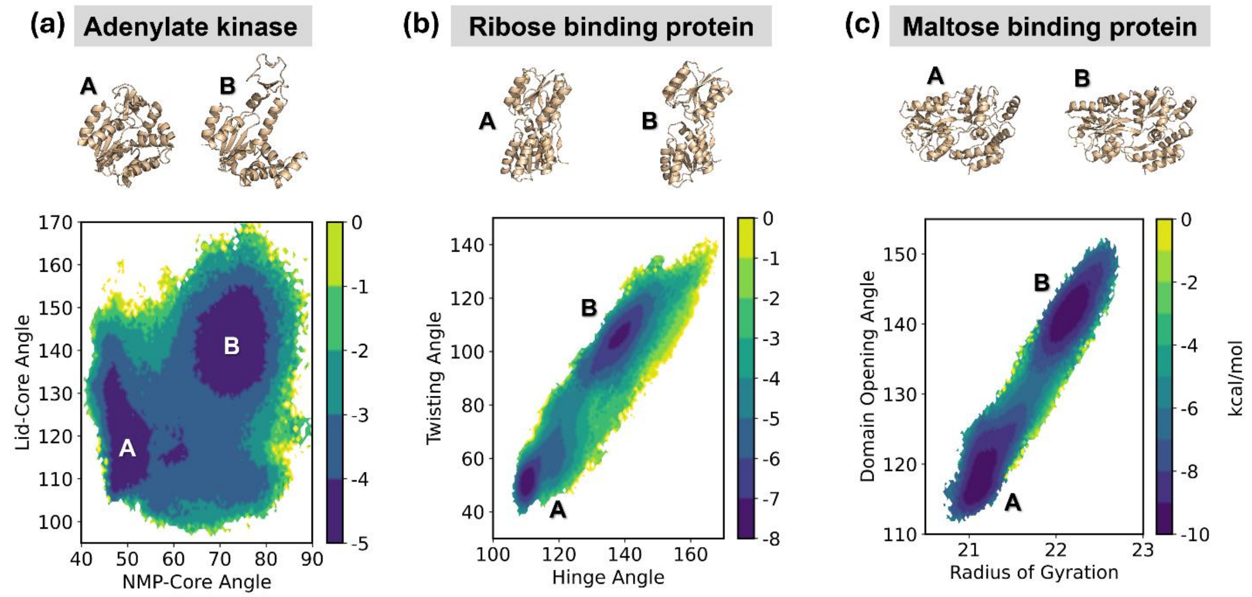


**Figure S4. Conventional WE simulations with a radial binning scheme plotted in common, physical coordinate subspaces for three medium-sized showcase protein systems, (a) adenylate kinase, (b) ribose-binding protein, and (c) maltose-binding protein.** The distributions shown are the combined result of six independent WE simulations, three for the A to B direction and three for the B to A direction, with WE weights normalized across the six independent simulations.
